# Rgs12 enhances osteoclastogenesis by suppressing Nrf2 activity and promoting the formation of reactive oxygen species

**DOI:** 10.1101/470542

**Authors:** Andrew YH Ng, Ziqing Li, Megan M Jones, Chengjian Tu, Merry J Oursler, Jun Qu, Shuying Yang

## Abstract

The Regulator of G-protein Signaling 12 (Rgs12) is important for osteoclast (OC) differentiation, and its deletion in vivo protected mice against pathological bone loss. To characterize its mechanism in osteoclastogenesis, we selectively deleted Rgs12 in OC precursors using the LysM-Cre transgenic line or overexpressed the gene in RAW264.7 cells. Rgs12 deletion led to increased bone mass with decreased OC numbers, whereas its overexpression increased OC number and size. Proteomics analysis of Rgs12-deficient OCs identified an upregulation of antioxidant enzymes under the transcriptional regulation of Nrf2, the master regulator of oxidative stress. We confirmed an increase of Nrf2 activity and impaired production in Rgs12-deficient cells. Conversely, Rgs12 overexpression suppressed Nrf2 through a mechanism dependent on the 26S proteasome, and promoted RANKL-induced phosphorylation of ERK1/2 and NFκB, which was abrogated by antioxidant treatment. We therefore identified a novel role of Rgs12 in regulating Nrf2, thereby controlling cellular redox state and OC differentiation.

## INTRODUCTION

Osteoporosis is a pervasive disorder characterized by skeletal fragility and microarchitectural deterioration that predisposes individuals to bone fractures. The disease has a significant global impact, affecting an estimated 200 million people worldwide and exerts a heavy economic burden—the affected are projected to rise by approximately 50% within the next 10 years (1). Therefore, understanding the pathogenesis of osteoporosis is an urgent matter to develop better treatments for this debilitating disease.

Bone remodeling is carried out by the coordinated actions of the bone-forming osteoblasts (OBs) and the bone-resorbing osteoclasts (OCs). Disorders of skeletal deficiency, such as osteoporosis, are typically characterized by enhanced osteoclastic bone resorption relative to bone formation, thereby resulting in net bone loss. Although significant progress has been made in OC behavior, it remains largely unknown about the regulatory mechanism that drives OC differentiation, which restricts the effectiveness of current treatments.

Regulators of G-protein Signaling (RGS) are a family of proteins comprised of more than 30 proteins that share a conserved RGS domain and play a classical role in attenuating G protein-coupled receptor (GPCR) signaling through its GTPase-accelerating protein (GAP) activity to inactivate the Gα subunit (2, 3). The RGS proteins are multifunctional proteins which hold vital cellular processes, including cell differentiation. Rgs12 is unique in that it is the largest protein in its family. In addition to the RGS domain, contains a PSD-95/Dlg/ZO1 (PDZ) domain, a phosphotyrosine-binding (PTB) domain, a tandem ras-binding domain (RBD1/2), and a GoLoco interaction motif. The multi-domain architecture of Rgs12 suggests a role as a scaffolding protein in complexes where multiple signaling pathways might converge (4-8).

Reactive oxygen species (ROS) are produced as a normal byproduct of cellular metabolism (9). Recent studies clearly show that RANKL-induced reactive oxygen species (ROS) are indispensable for OC differentiation (9-12). ROS at high levels induce oxidative stress, which if left unchecked becomes deleterious to cell. At low concentrations, however, ROS have been shown to participate in signaling events in OCs, including the RANKL-dependent activation of mitogen-activated protein kinases (MAPKs), phospholipase C gamma (PLCγ), nuclear factor kappa B (NFκB), and [Ca^2+^] oscillations; all of which contribute to the activation of nuclear factor of T-cells (NFAT), the master regulator of OC differentiation. Furthermore, multiple lines of evidence have consistently shown that suppression of ROS by various means inhibits OC differentiation (10-12). In particular, RANKL-dependent activation of PLCγ, [Ca^2+^] oscillations, and NFAT were abrogated when the OC precursors were treated with the antioxidant N-acetylcysteine (NAC) (11). Furthermore, our previous studies have that Rgs12 silencing could inhibit PLCγ activation, [Ca^2+^] oscillations, and the expression of NFATc1 and its downstream factors (13). Hence, these findings led us to hypothesize that Rgs12 may play a role in regulating the cellular redox state, thereby controlling OC differentiation.

Using an *in vivo* Rgs12 conditional knockout mouse model, we found that RANKL-dependent ROS was suppressed in Rgs12-deficient bone marrow macrophages (BMMs). Additionally, Rgs12-deficient cells have elevated levels of phase II detoxification and antioxidant enzymes. We further identified that Rgs12 deficiency increased the activation of Nrf2, the master transcription factor responsible for the expression of antioxidant proteins. Conversely, Rgs12 overexpression could suppress Nrf2 and promote osteoclastogenesis. Our results therefore demonstrate a novel function of Rgs12 in regulating ROS during OC differentiation, likely by suppressing Nrf2 to inhibit the expression of antioxidant proteins. These findings contribute to a better understanding of ROS regulation in OCs and future treatment strategies for diseases of excessive bone loss such as osteoporosis.

## RESULTS

### Targeted deletion of Rgs12 in myeloid cells increased trabecular bone mass

To assess the role of Rgs12 on OC differentiation and bone remodeling *in vivo*, we generated a conditional gene knockout mouse model by crossing Rgs12^flox/flox^ mice with LysM-Cre transgenic mice (LysM;Rgs12^fl/fl^). The LysM promoter-driven Cre expression targets the gene deletion to cells of the myeloid lineage, including monocytes/macrophages (14, 15). The Cre-lox-mediated deletion of the *Rgs12* gene was confirmed by PCR amplification of spleen genomic DNA (Fig. 1A), and qPCR to measure *Rgs12* transcripts in isolated BMMs (Fig. 1B) thereby confirming our mouse *Rgs12* conditional gene knockout model.

**Figure 1.**
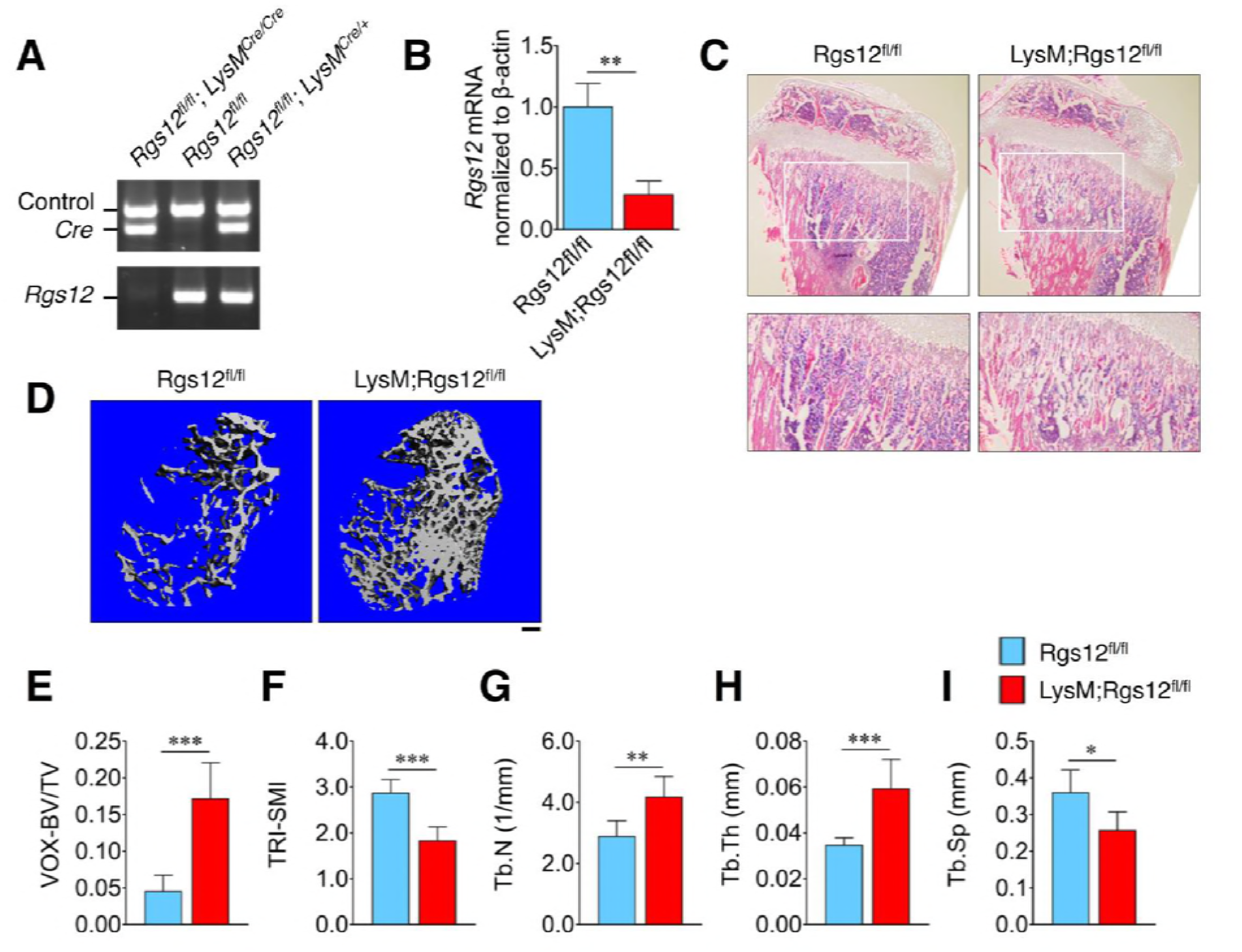
Rgs12-deficient mice exhibit increased trabecular bone mass. (A) PCR of splenic genomic DNA amplified the deletion allele of Rgs12 in LysM;Rgs12^fl/fl^ mice. (B) qPCR analysis of Rgs12 mRNA levels normalized to β-actin in BMMs obtained from Rgs12^fl/fl^ and LysM;Rgs12^fl/fl^ mice. Results are means ± SD (*n*=3, \*\**p*<0.01). (C) Hematoxylin and eosin staining of the proximal tibias and (D) Representative 3D micro-computed tomographay (micro-CT) images of the trabecular bone of femurs obtained from 8-weeks-old Rgs12^fl/fl^ and LysM;Rgs12^fl/fl^ mice. Scale bar on (D) = 200 μm. (E-I). Quantitative micro-CT measurements of femur bone morphology and microarchitecture. Results are means ± SD (*n*=4; \**p*<0.05, \*\**p*<0.01, \*\*\**p*<0.001). VOX-BV/TV, bone volume to tissue volume (voxel count); TRI-SMI, structure model index; Tb.Th, trabecular thickness; Tb.N, trabecular number; Tb.Sp, trabecular separation.

LysM;Rgs12^fl/fl^ mice exhibited an increase in trabecular bone mass, evident in the H&E-stained bone sections of the proximal tibia (Fig. 1C) and micro-computed tomography (micro-CT) analysis of the femoral trabecular bone morphology and microarchitecture (Fig. 1D and E). Quantitative micro-CT measurements further revealed significant increases in both trabecular number (Tb.N) and thickness (Tb.Th), and reduced trabecular separation (Tb.Sp) (Fig. 1G-I). Our results therefore demonstrate the importance of Rgs12 in bone remodeling.

### Rgs12 promotes osteoclast formation

To assess the role of Rgs12 in osteoclastogenesis, OC precursors isolated from LysM;Rgs12^fl/fl^ mice were differentiated using M-CSF and RANKL. While control BMMs differentiated into large, TRAP^+^ multinucleated OCs, Rgs12-deficient precursor cells showed a reduction in the number of OCs containing 6-9 nuclei and 10+ nuclei, which were also visibly smaller (Fig. 2A and B). Complementing our Rgs12 knockout model, we generated an Rgs12 overexpression OC model in which RAW264.7 cells were stably-transfected with a vector carrying a recombinant N-terminus FLAG-tagged Rgs12 gene (Flag-Rgs12). Rgs12 overexpression in RAW264.7 cells was confirmed by western blotting (Fig. 2C). Using this cell model, we next determined whether Rgs12 overexpression could promote osteoclastogenesis (Fig 3D). Converse to our findings in Rgs12 knockout cells, we found that overexpression of Rgs12 in RAW264.7 cells led to an increased number of OCs with 10+ nuclei (Fig. 3E). We also observed significantly decreased numbers of smaller OCs containing 3-5 and 6-9 nuclei in Rgs12 overexpressing cells (Fig. 2E), presumably because most of the smaller OCs have fused to form large OCs containing 10+ nuclei. In a study investigating the relationship between osteoclast size and state of resorptive activity, a greater proportion of large osteoclasts were active whereas non-resorbing osteoclasts were on average smaller (16). Additionally, quantification of the mean areas of OCs with 10+ nuclei revealed that Rgs12-overexpressing OCs were larger as compared to empty vector-transfected controls (Fig. 2F). Our findings consequently demonstrate the importance of Rgs12 in promoting OC formation, which is consistent with the osteopetrotic phenotype observed in Rgs12-deficient mouse model.

**Figure 2.**
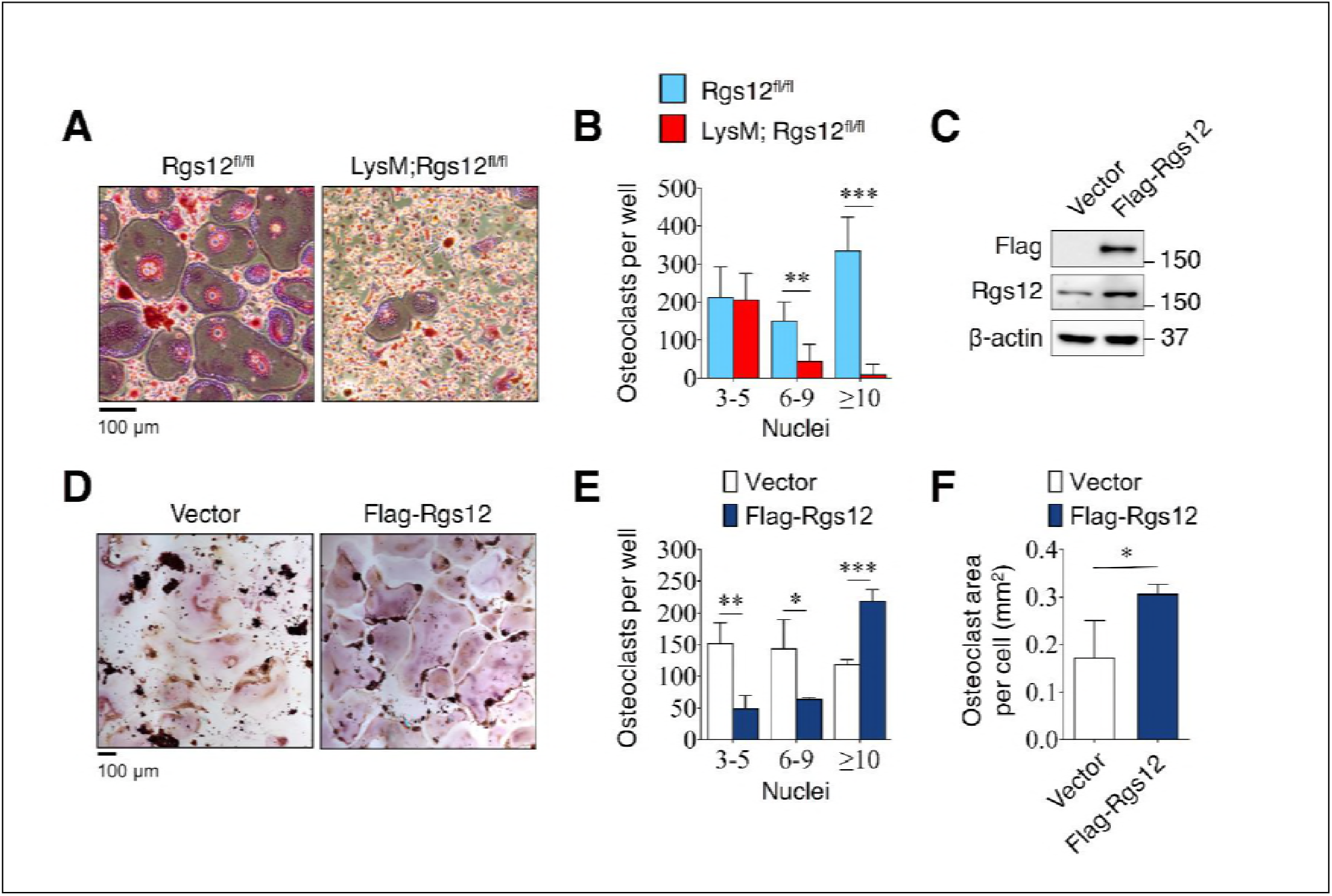
Rgs12 is essential for osteoclast differentiation. (A) TRAP-stained osteoclasts differentiated from BMMs isolated from Rgs12^fl/fl^ and LysM;Rgs12^fl/fl^ mice. (B) Number of TRAP¬positive and multinucleated (≥3 nuclei) osteoclasts from Rgs12^fl/fl^ and LysM;Rgs12^fl/fl^ BMMs (*n*=4, \**P*<0.05, \*\**P*<0.01, \*\*\**P*<0.001). (C) Immunoblot to verify Rgs12 overexpression in RAW264.7 cells transfected with a vector carrying a recombinant N-terminus FLAG-tagged Rgs12 gene (Flag-Rgs12). RAW264.7 cells transfected with the empty vector was used as a negative control. (D) TRAP-stained osteoclasts derived from RAW264.7 cells transfected with an empty vector or Flag-Rgs12. (E) Number of TRAP-positive and multinucleated (≥3 nuclei) osteoclasts from vector- and Flag-Rgs12-transfected RAW264.7 cells (*n*=3, \**P*<0.05, \*\**P*<0.01, *** *P*<0.001). (F) OC size was estimated by quantifying the surface area of osteoclasts containing 10+ nuclei, which was normalized to the number of osteoclasts with 10+ nuclei (*n*=3, \**P*<0.05). TRAP, tartrate-resistant acid phosphatase.

**Figure 3.**
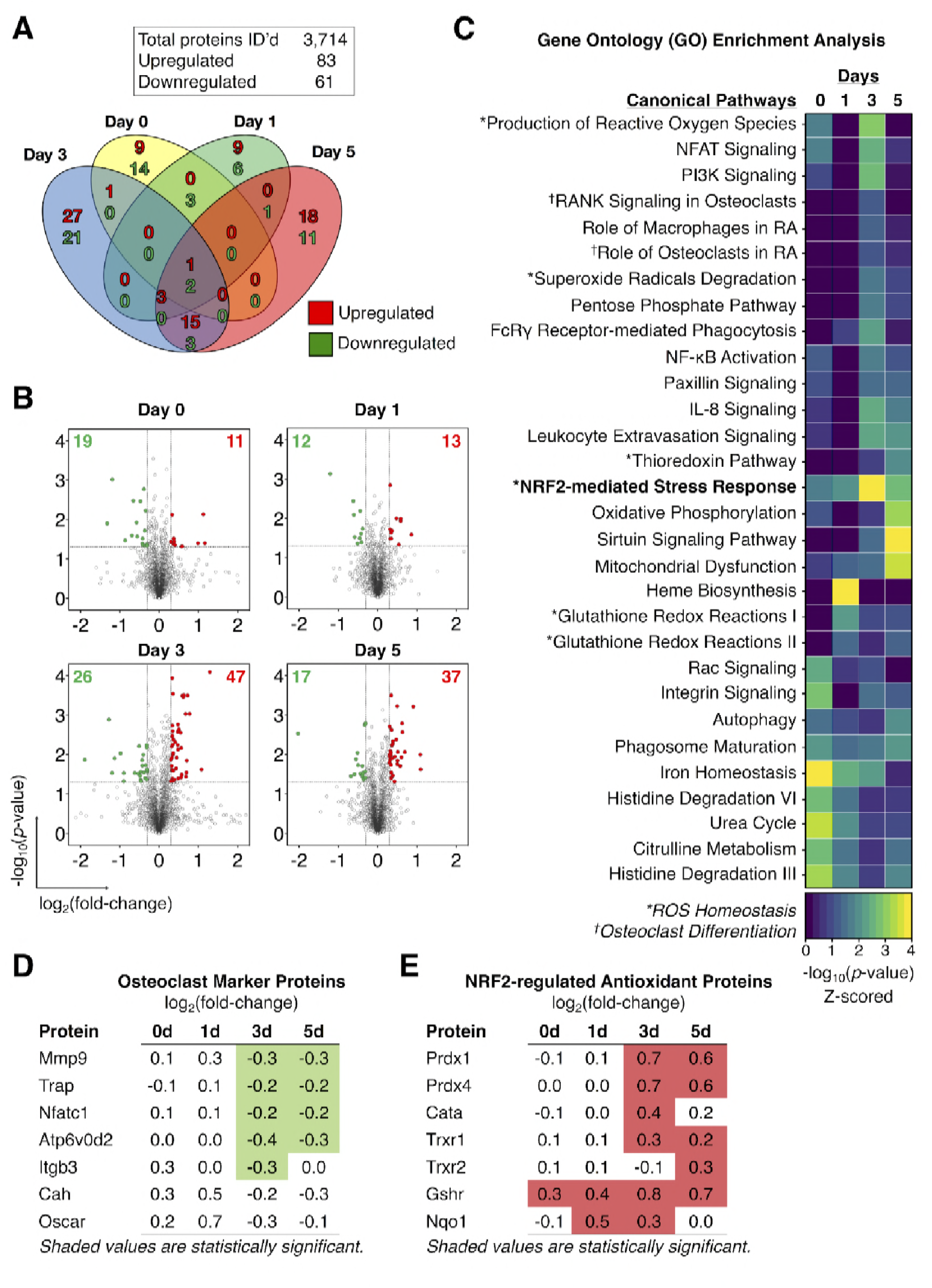
Proteomics analysis identified an increased expression of Nrf2-dependent antioxidant proteins in Rgs12-deficient osteoclast precursors. (A) Venn diagram summarizing the distribution of proteins that were significantly altered in LysM;Rgs12^fl/fl^ BMMs as compared to Rgs12^fl/fl^ BMMs at 0, 1, 3, and 5 days of OC differentiation. (B) Volcano plots depicting protein expression changes in LysM;Rgs12^fl/fl^ BMMs as compared to Rgs12^fl/fl^ BMMs. Optimized cutoff thresholds for significantly altered proteins was set at 1.3 log_2_-tranformed ratios and *p*-value < 0.05. Data are means ± SD. Student’s *t* test was performed to compare Rgs12^fl/fl^ and LysM;Rgs12^fl/fl^ BMMs at each time point (*n* = 3). (C) Gene ontology (GO) enrichment analysis to identify canonical pathways corresponding to the significantly altered proteins. For visualization purposes, the color intensity in the heat map diagram indicates the significance of GO term enrichment, presented as -log10(*p*-value). Hierarchical clustering analysis was used to group GO terms based on the *p*-value of enrichment. (D, E) The expression of OC marker proteins and Nrf2-regulated antioxidant proteins in LysM;Rgs12^fl/fl^ versus Rgs12^fl/fl^ BMMs. Mmp9, metalloproteinase-9; Trap, tartrate-resistant acid phosphatase; Nfatc1, nuclear factor of activated T cells, cytoplasmic 1; Atp6v0d2, ATPase H+ transporting V0 subunit D2; Itgb3, integrin β3; Prdx, peroxiredoxin; Cata, catalase; Trxr, thioredoxin; Gshr, glutathione reductase; Nqo1, NAD(P)H dehydrogenase quinone 1.

### Rgs12-defcient osteoclast precursors show an increased expression of Nrf2-dependent antioxidant proteins

To uncover the role of Rgs12 in OC differentiation, we employed the *IonStar* liquid chromatography tandem mass spectrometry (LC-MS/MS)-based quantitative proteomics strategy (17) to profile the temporal dynamics in the global protein levels in Rgs12^fl/fl^ and LysM;Rgs12^fl/fl^ BMMs at 0, 1, 3, and 5 days of OC differentiation (Fig. 4). Proteomics analysis identified 3,714 quantifiable proteins that are present in all samples (no missing data), using a highly stringent identification criteria of ≥2 peptides per protein and 1% false discovery rate (Fig. 3A). Within this dataset, we identified 83 and 61 unique proteins that were significantly up- and downregulated, respectively, in LysM;Rgs12^fl/fl^ BMMs relative to Rgs12^fl/fl^ BMMs. Proteins were considered significantly altered if they exceeded the thresholds set at *p*<0.05 and >0.3 log_2-_transformed ratio (Fig. 3B). Most of the protein expression changes in Rgs12-deficient cells were captured at 3 and 5 days of OC differentiation (Fig. 3A). To determine the biological significance of these altered proteins, we performed gene ontology analysis to identify the canonical pathways involved (Fig. 3C). Processes related to OC differentiation (e.g. “NFAT Signaling”, “RANK Signaling in OCs”, and “Role of OCs in Rheumatoid Arthritis”) were enriched at 3 and 5 days of OC differentiation. Closer inspection showed that OC marker proteins including metalloproteinase-9 (Mmp9), tartrate-resistant acid phosphatase (Trap), ATPase H^+^ transporting V0 subunit D2 (Atp6v0d2), and integrin β3 (Itgb3) were significantly downregulated in LysM;Rgs12^fl/fl^ OCs (Fig. 3D). Additionally, the analysis revealed several biological functions related to ROS homeostasis that were impacted by Rgs12 deletion (e.g. “Production of ROS”, “Superoxide Radical Degradation”, and NRF2-mediated Stress Response”) (Fig. 3C). Inspection of the proteins involved in these pathways showed a significant upregulation of numerous Nrf2-dependent antioxidant enzymes responsible attenuating oxidative stress, including: peroxiredoxin 1/4 (Prdx1/4), thioredoxin 1/2 (Trxr1), glutathione reductase (Gshr), and NAD(P)H dehydrogenase quinone 1 (Nqo1) (Fig. 3E). Upstream regulator (transcription factor) analysis identified that the antioxidant enzymes upregulated in LysM;Rgs12^fl/fl^ OCs share the common upstream regulator Nrf2, a key transcription factor that regulates cellular redox balance through the expression of protective antioxidant and phase II detoxification proteins. (18, 19) (Fig S4A). Although the upstream regulator analysis predicted an upregulation of Nrf2 activity, the transcription factor itself was not detected by our proteomics analysis. Proteins of typically low abundance such as cytokines, signal regulatory molecules, and transcription factors tend to be “crowded out” during MS analysis by more highly abundant proteins such those proteins involved in glycolysis and purine metabolism, protein translation, and cytoskeletal components (20). Nonetheless, our proteomics-based discovery tool allowed us to generate the hypothesis that Nrf2 is aberrantly activated by Rgs12 deletion, causing excessive clearance of ROS by antioxidant enzymes and in turn disrupting OC differentiation.

**Figure 4.**
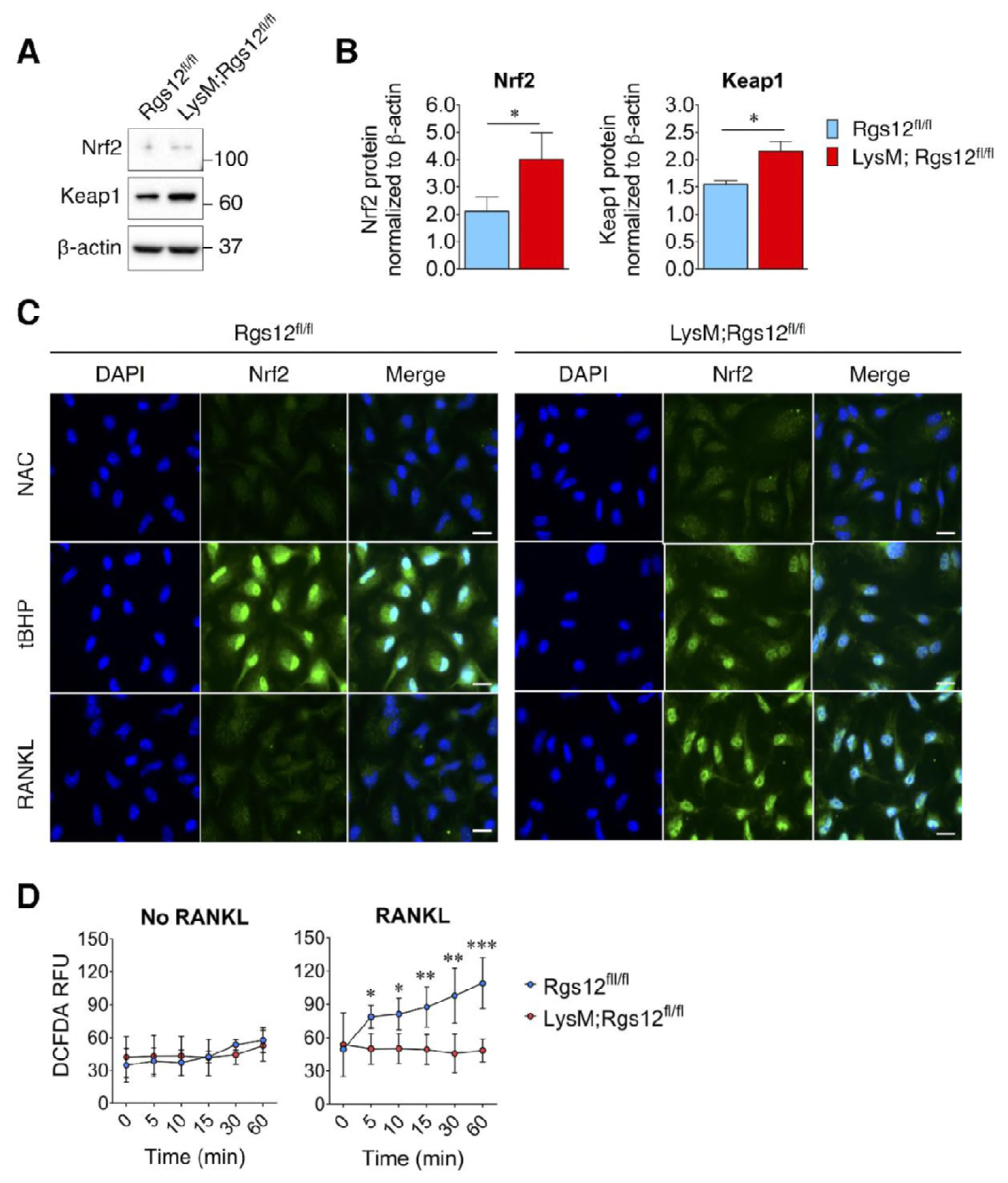
Increased Nrf2 activation and expression of antioxidant proteins in Rgs12-deficient osteoclast precursors. (A-F) Nrf2 immunofluorescence staining in Rgs12^fl/fl^ and LysM;Rgs12^fl/fl^ BMMs differentiated with M-CSF and RANKL for 72 h. (C, D). As a negative control for Nrf2 nuclear translocation, cells were treated with the antioxidant compound NAC (5 mM, 16 h) to suppress cellular ROS. (E, F) Conversely, as a positive control for Nrf2 nuclear translocation, cells were treated with the peroxidase tBHP (50 μM, 16 h) to induce oxidative stress. (G, H) Immunoblot of Nrf2 and Keapl protein levels in Rgs12^fl/fl^ and LysM;Rgs12^fl/fl^ BMMs treated with RANKL for 72 h. Densitometry analysis was performed on bands and normalized to β-actin (*n*=3, \**p*<0.05). (I) Induction of ROS levels in Rgs12^fl/fl^ and LysM;Rgs12^fl/fl^ BMMs differentiated for 72 h, kept in serum-free medium for 6 h, and stimulated with RANKL for the indicated times. ROS levels were measured using the DCFDA fluorescence method. Data are means ± SD (*n*=5, \**p*<0.05, \*\**p*<0.01, \*\*\**p*<0.001). DAPI, 4,6-diamidino-2-phenylindole; NAC, N acetylcysteine; tBHP, fe/f-butylhydroxyperoxide. ROS, reactive oxygen species. DCFDA, 2’,7’-dichlorofluorescin diacetate. RFU, relative fluorescence units.

### Deletion of Rgs12 elevated Nrf2/Keap1 expression and Nrf2 activity

Based on our proteomics analysis, we hypothesized that Rgs12 is needed to suppress Nrf2 activity and facilitate the formation of ROS, which has been previously shown to play a critical role in OC differentiation (10, 21). To test this hypothesis, we assessed Nrf2 activity and the expression of Nrf2 and Keap1 in Rgs12^fl/fl^ and LysM;Rgs12^fl/fl^ BMMs (Fig. 4). Western blotting of Nrf2 in day 3 OCs also showed increased levels of Nrf2 in LysM;Rgs12^fl/fl^ cells (Fig. 4A and B). Keap1, however, which is known to suppresses Nrf2 activity by facilitating its degradation via the proteasome pathway, was unexpectedly elevated in Rgs12-deficient cells. Furthermore, immunofluorescence staining of Nrf2 demonstrated increased nuclear translocation of the transcription factor in LysM;Rgs12^fl/fl^ cells (Fig. 4C, upper panel). The elimination of ROS with N-acetylcysteine (NAC), a precursor to the antioxidant glutathione, was able to completely suppress Nrf2 nuclear translocation in both Rgs12^fl/fl^ and LysM;Rgs12^fl/fl^ BMMs (Fig. 4C, middle panel). Conversely, induction of oxidative stress using the peroxide *tert*-buthylhydroxyperoxide (tBHP) potently induced Nrf2 nuclear translocation (Fig. 4C, bottom panel). To further test whether elevated Nrf2 activity in Rgs12-deficient OCs could result in reduced intracellular ROS levels, we detected intracellular ROS levels. As expected, the RANKL-dependent ROS induction observed in control cells was suppressed in LysM;Rgs12^fl/fl^ OCs (Fig. 4D). These findings demonstrate an abnormal upregulation of Nrf2 activity and expression in Rgs12-deficient cells, indicating that Rgs12 may be required to suppress Nrf2 to facilitate osteoclastogenesis.

### Rgs12-mediated suppression of Nrf2 activity is dependent on the proteasome degradation pathway

Under basal conditions (i.e. absence of cellular stress), Nrf2 remains inactive through its interaction with Keap1, which causes its continual ubiquitination and degradation via the proteasome pathway (22, 23). A variety of stress conditions can induce conformational changes in Keap1, thereby releasing Nrf2 from the ubiquitin-proteasome pathway, allowing it to accumulate and translocate into the nucleus (24, 25). To better understand the mechanism by which Rgs12 suppresses Nrf2 activity, we therefore first determined whether the ability of Rgs12 to suppress Nrf2 activity relies on this canonical mechanism (Fig. 5A). Given that Rgs12 deletion resulted in elevated Nrf2 expression and nuclear translocation, we first determined whether Rgs12 overexpression could exert the opposite effect (Fig. 5A and B). We measured Nrf2 protein levels in RAW264.7 cells stably transfected with the Rgs12-His or empty vector and found no difference when cells are at their basal state (i.e. uninduced). Stimulation of RAW264.7 cells with *tert*-buthylhydroquinone (tBHQ), which is known to directly bind Keap1 and attenuate its inhibitory effect on Nrf2 (26), caused a robust induction of Nrf2 protein levels in a dose-dependent manner (Fig. 5A and B). More importantly, RAW264.7 cells overexpressing Rgs12 showed a significant reduction of Nrf2 protein levels that resulted from Keap1 inhibition compared to those in the control cells. Moreover, the ability of Rgs12 to facilitate Nrf2 degradation despite the inhibition of Keap1 suggests that Rgs12 functions downstream of Keap1, either by controlling the ubiquitination or proteasomal degradation of Nrf2.

**Figure 5.**
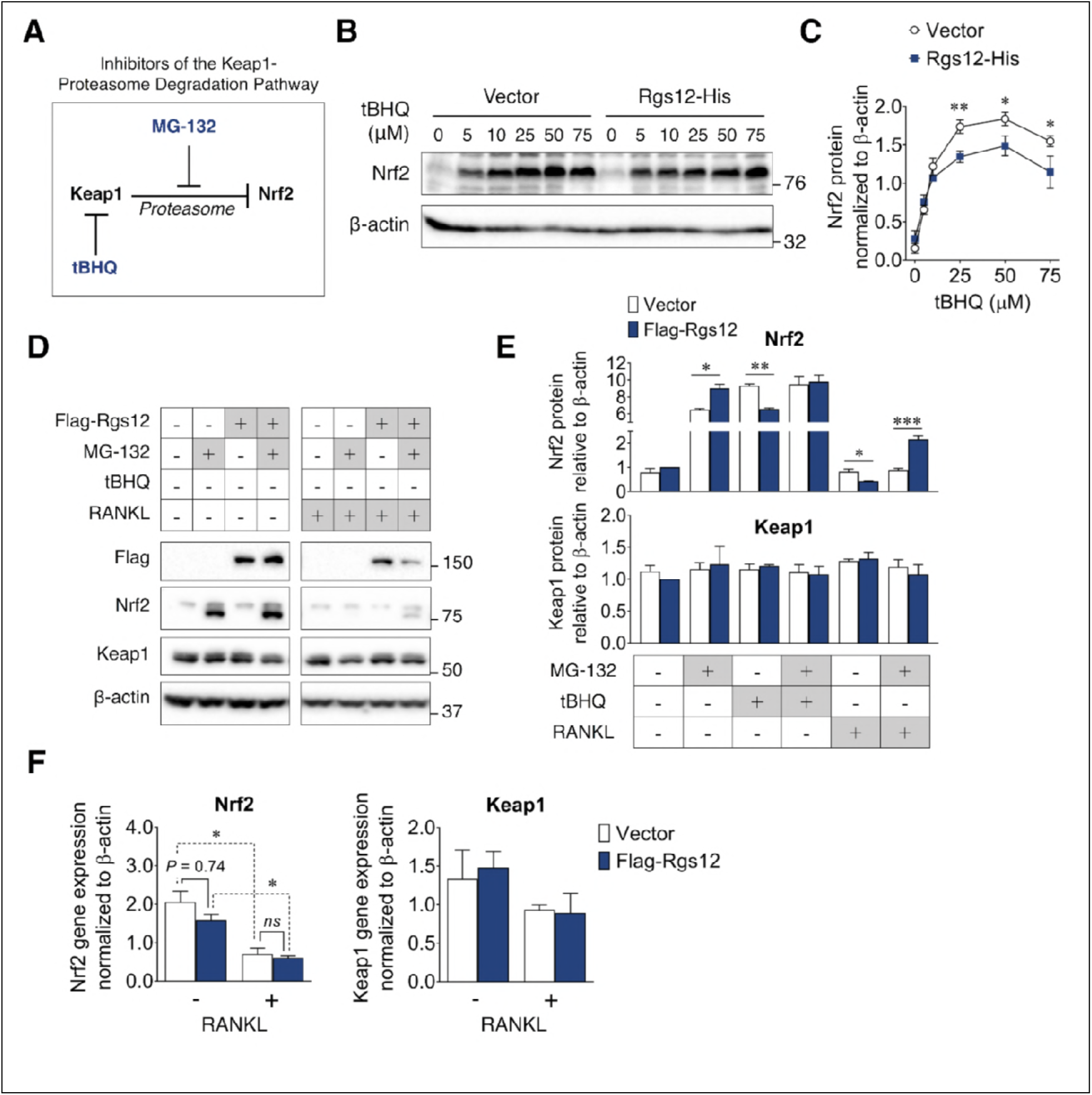
Suppression of Nrf2 protein levels by Rgs12 is dependent on the proteasome degradation pathway. (A) Diagram summarizing the inhibitors of the Keap1-proteasome axis to modulate Nrf2 protein levels. (B) RAW264.7 cells stably-transfected with Rgs12-His or empty vector treated with increasing doses of tBHQ. (C) Nrf2 and Keap1 protein levels were quantified by densitometry analysis and normalized to β-actin (*n*=3, \**p*<0.05, \*\**p*<0.01). (D) Western blot to detect Nrf2 and Keap1 in RAW264.7 cells stably-transfected with empty vector or Flag-Rgs12. RAW264.7 cells were treated with a combination of RANKL (100 ng/mL, 72 h) and the proteasome inhibitor MG-132 (25 μM, 4 h). (E) Nrf2 and Keap1 protein levels were quantified by densitometry analysis and normalized to β-actin (*n*=3, \**p*<0.05, \*\**p*<0.01, \*\*\**p*<0.001). (F) qPCR analysis of Nrf2 and Keap1 transcript levels in RAW264.7 cells transfected with Rgs12-His or empty vector. Data are means ± SD. Two-tailed *t* test was performed (*n*=3, \**P*<0.05). tBHQ, *tert-* butylhydroquinone.

Given the possibility that the reduction of Nrf2 levels in Rgs12 overexpression cells may be a result of increased Nrf2 degradation, we further tested whether inhibiting the proteasome, a step downstream of Keap1, could attenuate the ability of Rgs12 to facilitate Nrf2 degradation (Fig. 5D and E). Similar to tBHQ, preventing Nrf2 degradation using the proteasome inhibitor MG-132 caused Nrf2 protein to substantially accumulate (Fig. 5D, left panel). Interestingly, when Nrf2 protein levels were artificially induced, we observed the presence of a lower molecular weight band, which could correspond to a different post-translational modification state (e.g. unphosphorylated or non-ubiquitinated). Furthermore, we did not observe any changes in Keap1 protein levels. In the previous scenario wherein Rgs12 overexpression could still promote Nrf2 degradation in spite of tBHQ treatment, this was not the case when using MG-132. In fact, inhibiting the proteasome was able to reverse the ability of Rgs12 to promote Nrf2 degradation, indicating its requirement for the proteasome’s function. Repeating this experiment in RAW264.7 cells differentiated for 3 days with RANKL showed that Nrf2 levels were suppressed in OCs (Fig. 5D, right panel); likely due to reduced transcriptional activity, which corroborates with findings from previous studies (Fig. 5F) (21, 27). More importantly, Rgs12 overexpression could suppress Nrf2 protein levels, but inhibition of the proteasome using MG-132 reversed this effect (Fig. 5D, right panel). To confirm that Rgs12 inhibits Nrf2 through a post-translational mechanism, we measured Nrf2 transcript levels by qPCR and found no difference between wild-type or Rgs12-overexpressing cells (Fig. 5F). Overall, our data collectively indicate that Rgs12 suppresses Nrf2 activity by facilitating its degradation through the proteasome-dependent pathway.

### Rgs12-mediated activation of osteoclast MAPK and NFκB signaling is dependent on intracellular ROS

It was previously demonstrated that ROS could act as an intracellular signal mediator OC differentiation, and is required for the RANKL-dependent activation of p38 mitogen-activated protein kinase (MAPK), extracellular signal-regulated kinase (ERK), and NFκB (10, 28). Given our findings that Rgs12 could suppress the activity of Nrf2 and thereby promoting intracellular ROS, we hypothesized that Rgs12 could promote RANKL-dependent signaling, and that this effect would be abrogated by the addition of an antioxidant (Fig. 6A and B). As expected, RAW264.7 cells overexpressing Rgs12 demonstrated a more robust activation of ERK1/2 and NFκB phosphorylation but not p38 MAPK. Pretreating Rgs12 overexpressing cells with the antioxidant NAC diminished ERK1/2 activation, and almost completely abrogated NFκB activation. These results support the role of Rgs12 in promoting ROS that is important OC signaling, likely through the suppression of Nrf2 activity.

**Figure 6.**
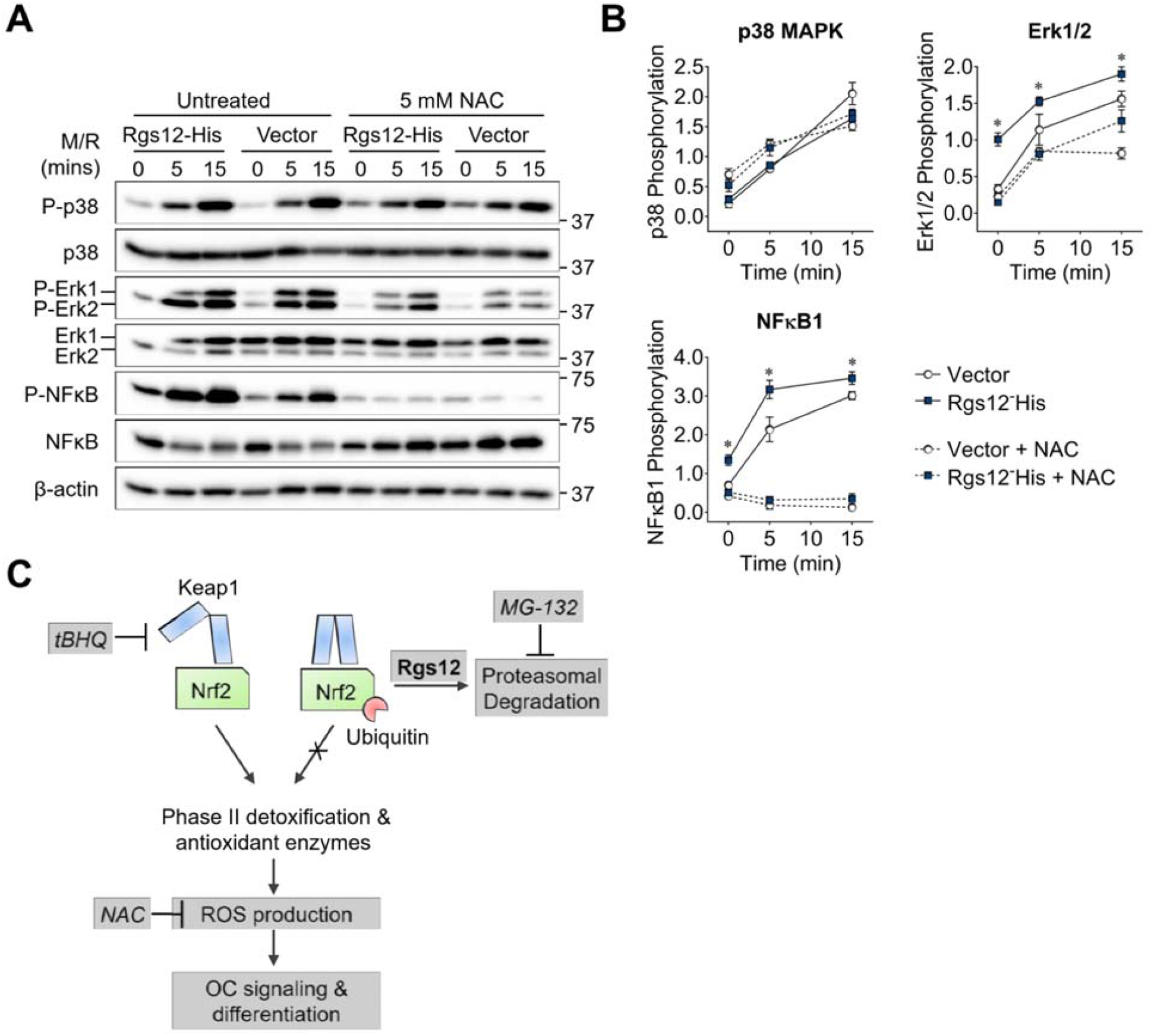
Rgs12-dependent activation of MAPK and NFkB was suppressed by antioxidants. (A)Western blot detected phosphorylated or total p38, NFκB, and Erk1/2 in transfected RAW264.7 cells induced with RANKL (200 ng/mL) and M-CSF (100 ng/mL) for the indicated times. Cells were pretreated with NAC (5 mM, 4 h) to suppress cellular reactive oxygen species. (B) Band density was quantified by ImageJ and phosphorylated and unphosphorylation/total protein levels were normalized to β-actin. Relative phosp 1 horylation are presented as the ratio between the phosphorylated normalized to the nonphosphorylated/total protein. Two-tailed *t* tests were used to compare vector and Rgs12-His groups (*n*=3, \**P*<0.05). M/R, M-CSF and RANKL. NAC, N-acetylcysteine. (C) Model of the role of Rgs12 in suppressing Nrf2 to promote ROS and OC differentiation.

## DISCUSSION

The importance of ROS in osteoclasts has been underlined by the growing body of evidence that ROS increased with aging or during inflammation can stimulate bone resorption and exacerbate bone loss (9). Targeting ROS in diseases of excess bone resorption such as osteoporosis could therefore be an important therapeutic strategy. Additionally, an important mechanism of cellular ROS clearance relies on the Keap1-Nrf2 pathway which is very well characterized, especially in the context of cancer biology (29). However, the upstream signaling molecules that could regulate the Keap1-Nrf2 axis in OCs remains unknown. Targeting this gap in knowledge, our study uncovered a novel role of the signaling protein Rgs12 in regulating Nrf2, thereby controlling cellular redox state and OC differentiation (Fig. 6C).

In this study, we demonstrated the essential role of Rgs12 in OC differentiation such that Rgs12 knockout mice exhibited increased bone mass (Fig. 1) and OC precursors isolated from these mice showed reduced OC differentiation (Fig. 2). On the contrary, overexpressing Rgs12 in RAW264.7 cells significantly promoted OC formation and increased the size of resultant OCs. However, the mechanism by which Rgs12 regulates OC differentiation remains unclear.

Proteomics is a powerful tool that has led to numerous discoveries of proteins and biological processes that drive OC differentiation (30). Notably, this technique was recently used to map the podosome proteome which helped to advance our understanding of determinants in the macrophage multinucleation process (31, 32), and how metabolism and energy is redirected towards bone resorption in OCs (33). Proteomics can therefore provide a broad yet informative overview of the systemic changes in the differentiating OC. To discover the cellular function of Rgs12 in OCs, we employed a robust and high-throughput quantitative proteomics approach to characterize the global protein changes of OCs derived from Rgs12^fl/fl^ and LysM;Rgs12^fl/fl^ BMMs (Fig. 3). Interestingly, the analysis identified the upregulation of a collection of antioxidant enzymes that are transcriptionally regulated by the antioxidant response element (ARE) in the promoter region, which is activated by the transcription factor Nrf2 (34). Based on this evidence, we further investigated the role of Rgs12 in Nrf2 signaling. We found that Nrf2 protein levels and nuclear translocation were increased in OC precursors in which Rgs12 was deleted (Fig. 4). Because our data showed that Rgs12 deficiency upregulated Nrf2, we expected that Keap1 levels should be reduced in order to facilitate the increased Nrf2 activity. On the contrary, Keap1 levels were upregulated in Rgs12-deficient cells, suggesting that the Nrf2 upregulation may be independent of the Keap1. We speculate that Rgs12-deficient cells may be overcompensating Keap1 expression in order to rein back the increased Nrf2 activity. Nevertheless, consistent with the upregulation of Nrf2 and its corresponding antioxidant enzymes, the RANKL-dependent induction of ROS was attenuated in Rgs12-deficient cells. Taking an opposite approach, we demonstrated that Rgs12 overexpression in OC precursors could also enhance RANKL-mediated activation of ERK and NFκB, which is known to be dependent on ROS (Fig. 7). Inhibition of intracellular ROS blocked the effect of Rgs12 overexpression, indicating that Rgs12 promotes RANKL-dependent signaling by facilitating ROS production. Overall, the data collectively demonstrate that Rgs12 promotes osteoclastogenesis by facilitating ROS generation through the suppression of Nrf2 and its target antioxidant genes.

Nrf2 is constitutively expressed but its activity is inhibited through its interaction with Keap1. Under basal conditions, Nrf2 is restricted to the cytoplasm where it is continually depleted through the proteosomal degradation pathway. When bound to Nrf2, Keap1 recruits the Cul3-dependent E3 ubiquitin ligase complex, which ubiquitinates and targets Nrf2 for degradation by the 26S proteasome (23, 24, 35, 36). The Keap1 protein containing multiple reactive cysteine residues that serve as redox sensors (26) Stressor conditions including oxidative stress causes the electrophilic modification of Keap1, inducing conformational changes which cause the protein to dissociate from Nrf2 and allow the transcription factor to translocate into the nucleus (22). We therefore determined how Rgs12 regulates this well-defined mechanism (Fig. 5). tBHQ is a selective inhibitor of Keap1 activity by covalently binding the protein’s reactive thiols and could activate Nrf2 and its downstream proteins in RAW264.7 cells (26). Furthermore, tBHQ inhibited OC differentiation via the upregulation of heme oxygenase-1, a Nrf2-dependent antioxidant enzyme (Yamaguchi et al., 2014). In our study, we determined whether Rgs12 could suppress the tBHQ-dependent upregulation of Nrf2 (Fig. 5B). We reasoned that if Rgs12 relies on a Keap1-dependent mechanism, then the inhibition of Keap1 by tBHQ should prevent the ability of Rgs12 to suppress Nrf2. However, we observed that Rgs12 was still able to suppress Nrf2 despite blocking Keap1 activity, indicating that Rgs12 functions downstream of Keap1. Following Keap1-mediated ubiquitination of Nrf2, the targeted protein is degraded by the proteasome. Again, we reasoned that if Rgs12 is dependent on the proteasomal degradation pathway, then inhibition of this pathway using MG-132 should prevent Rgs12-mediated suppression of Nrf2. Indeed, we found that Inhibiting the proteasome was able to reverse the Rgs12-mediated degradation of Nrf2, which places Rgs12 in between Keap1 and the proteasome in the Nrf2 degradation pathway (Fig. 5D and E). Thus, Rgs12 could either regulate the Cul3-dependent E3 ubiquitin ligase complex to facilitate the ubiquitination of Nrf2, or directly control proteasome activity. It is interesting to note that NFκB activation is also dependent on the proteasomal degradation of inhibitor of κB (IκB), which otherwise sequesters NFκB to the cytoplasm (37). If Rgs12 could modulate proteasome activity, it is possible that Rgs12 could directly promote NFκB by facilitating the degradation of IκB. However, the fact that antioxidant treatment to suppress ROS could almost completely block the phosphorylation of NFκB points to an important role of ROS, and not just the proteasomal degradation of IκB in NFκB activation (Fig. 6). This potential crosstalk between the NFκB and Nrf2 pathways will need to be evaluated in future studies.

In conclusion, our study points to a novel role of Rgs12 in OC redox biology, thus forming the molecular basis for developing therapies for osteoporosis and other diseases of bone loss. Additionally, we found a new factor that could modulate the Nrf2-Keap1 pathway, which is important within the context of ROS biology.

## METHODS

### Generation of Rgs12 Conditional Knockout Mice

Rgs12^fl/fl^ mice were crossed with LysM-Cre transgenic mice to generate Rgs12 conditional knockout mice specific to the myeloid lineage (LysM;Rgs12^fl/fl^). The methodology for generating Rgs12^fl/fl^ and LysM-Cre mice and genotyping are previously described (15, 38, 39). Mice used for experiments were 6-8-weeks-old. All animal studies were approved by the University at Buffalo Institutional Animal Care and Use Committee (IACUC).

### Histology and quantitative micro-CT measurements

Mouse femurs were excised, fixed for 24 h in 10% natural buffered formalin, and decalcified in 10% EDTA for 1-2 weeks at 4 °C. The samples were embedded in paraffin and sectioned at 5 µm and stained with H&E. A quantitative analysis of the gross bone morphology and microarchitecture was performed using a micro-CT system (USDA Grand Forks Human Nutrition Research Center, Grand Forks, ND, USA). Fixed femur from Rgs12 control and mutant mice were analyzed and 3D reconstruction was used to determine bone volume to tissue volume (BV/TV), structure model index (SMI), trabecular thickness (Tb.Th, μm), trabecular number (Tb.N, /mm), and trabecular separation (Tb.Sp, μm).

### Generation of Rgs12 expression vectors

Full length Rgs12 (Accession: NM_173402.2) cDNA was cloned into the p3XFLAG-myc-CMV-26 expression vector (Sigma-Aldrich, St. Louis, MO, USA). Briefly, *Hind*III sites were incorporated into both termini of the Rgs12 cDNA using restriction-site-generating PCR, and the restriction sites were used to insert the Rgs12 sequence into the expression vector containing an N-terminus FLAG tag sequence (Flag-Rgs12). The primer walking method was used to validate the correct directionality of the insert. Additionally, a vector expressing C-terminus His-tagged Rgs12 (Rgs12-His) was generated by subcloning the Rgs12 cDNA into the pcDNA3.1(+)-c-His vector (Genscript, Piscataway, NJ, USA). *Hind*III and *Eco*RV sites were introduced by PCR and the restriction sites were used to insert Rgs12 into the His vector. All vector constructs were confirmed by DNA sequencing (Eurofins Genomics, Louisville, KY, USA).

### Stable Transfection

RAW264.7 cells were seeded at 2 × 10^6^ cells per 6-well and transfected using FuGENE HD reagent (Promega, Madison, WI, USA) according to manufacturer’s instruments at a 1:3 DNA to transfection reagent ratio. After 48 hours post-transfection, cells were treated with 0.4 mg/mL geneticin (G418, Thermo Fisher Scientific, Waltham, MA, USA) for 2 weeks until antibiotic-resistant colonies are formed. Stably transfected cells were thereafter maintained in media containing 0.4 mg/mL G418.

### *In Vitro* Osteoclastogenesis and TRAP staining

The vector encoding the recombinant mRANKL-His (K158-D316) construct and a modified *E. Coli* strain Origami B(DE3) cells (EMD Millipore, Billercica, MA, USA) co-expressing chaperone proteins that was used to express the recombinant RANKL were gifts from Dr. Ding Xu (University at Buffalo, School of Dental Medicine, Buffalo, NY). The protocol for expressing and purifying mRANKL-His was described previously (40). Endotoxins were removed using the Pierce High Capacity Endotoxin Removal Resin (Thermo Fisher Scientific, Waltham, MA, USA). The M-CSF-producing cell line CMG14-12 was a gift from Dr. Sunao Takeshita (National Center for Geriatrics and Gerontology, Obu, Japan). M-CSF production and bioassay were performed as previously described (41).

BMMs were obtained from the tibia and femur of 8-weeks-old C57BL/6J mice as described previously (13). BMMs were seeded at 2 × 10^6^ cells per 24-well and stimulated with 100 ng/mL RANKL and 20 ng/mL M-CSF for 5 days to generate mature OCs. RAW264.7 cells were seeded at 1.35 × 10^4^ cells per 24-well and stimulated with 100 ng/mL RANKL for 5 days. Prior to fixing and staining, RAW264.7-derived OCs were rinsed thoroughly with PBS to remove mononuclear cells that tend to obscure OCs during imaging. TRAP staining was performed using the acid phosphatase, leukocyte (TRAP) kit (Sigma-Aldrich, St. Louis, MO, USA). Cells were imaged using the Cytation 5 Cell Imaging Multi-Mode Reader (BioTek, Winooski, VT, USA) using the montage function. Osteoclasts were quantified by counting the number of TRAP^+^, multinucleated cells (MNCs, ≥3 nuclei/cell) per well. Average osteoclast area was determined by measuring total TRAP^+^ area using the ImageJ software (US National Institute of Health, Bethesda, MA, USA) and dividing the value by total osteoclast number. (41)

### Reverse Transcription and Quantitative PCR

Total RNA was isolated from cultured BMMs and OCs using Trizol reagent (Invitrogen, Carlsbad, CA, USA) following manufacturer’s instructions. cDNA was reverse transcribed from 2 μg total RNA using the RNA to cDNA EcoDry Premix kit (Clontech, Palo Alto, CA, USA). Primers were designed using Primer-BLAST (42) and obtained from IDT (Integrated DNA Technologies, San Diego, CA, USA). Rgs12 (F: 5’-AAGATCCATTCCCTAGTGACC-3’, R: 5’-ACCTCCACTTTCCCACCCTG-3’, 587 bp), Nrf2 (F: 5’-GCCCACATTCCCAAACAAGAT-3’, R: 5’-CCAGAGAGCTATTGAGGGACTG-3’, 172 bp), Keap1 (F: 5’-TGCCCCTGTGGTCAAAGTG-3’, R: 5’-GGTTCGGTTACCGTCCTGC-3’, 104 bp), β-actin (F: 5’-CTAGGCACCAGGGTGTGAT-3’, R: 5’-TGCCAGATCTTCTCCATG TC-3’, 148 bp). qPCR was performed using the 2x SYBR Green qPCR Master Mix following manufacturer’s instructions (Bimake, Houston, TX, USA). All reactions were performed in triplicate and normalized to the housekeeping gene β-actin. Data analysis was performed using the CTX Maestro software (Bio-Rad, Hercules, CA, USA).

### Fluorescent actin cytoskeleton staining

Cells were starved in medium containing 1% fetal bovine serum (FBS) and 5 ng/mL M-CSF for 16 hours then 0% FBS and 0 ng/mL M-CSF for 24 hours. Cytoskeletal reorganization was induced by introducing 50 ng/mL M-CSF in serum-free medium for 5 minutes. Cells were fixed and the actin cytoskeleton was visualized by rhodamine phalloidin staining.

### Rac1-GTP pulldown assay

The Rac1-GTP pulldown assay was performed following manufacturer instructions in the Rac1 activation assay kit (Cytoskeleton, Denver, CO, USA).

### ROS measurement

To measure ROS production, BMMs were seeded into black, glass-bottom 96-well plates and cultured with M-CSF for 48 hours until confluence. Cells were loaded with 20 μM 2’7’-dichlorofluorescein diacetate (DCFDA, Sigma) at 37 °C for 30 minutes and washed using PBS. The cells were swapped into complete phenol red-free MEM (Gibco) containing RANKL/M-CSF. Fluorescence intensity was measured using the Cytation 5 plate reader (BioTek) with excitation wavelength at 488 nm and emission wavelength at 535 nm. Background signals (cells not loaded with DCF-DA) were subtracted. Experiments were carried out in quintuplicate wells.

### Protein Extraction and Precipitation/ On-Pellet Digestion

Cells were harvested using ice-cold lysis buffer (50 mM Tris-formic acid, 150 mM NaCl, 0.5% sodium deoxycholate, 1% SDS, 2% NP-40, pH 8.0) with protease inhibitor (cOmplete, Mini, EDTA-free; Roche, Mannheim, Germany). Samples were prepared for MS analysis using an established method (17, 43).

### Liquid Chromatography-Tandem Mass Spectrometry Analysis

The “IonStar” LC-MS experimental pipeline was developed and optimized in a previous study (17, 43). A stringent set of criteria including a low peptide and protein false discovery rate (FDR) of < 1% and ≥2 peptides per protein was used for protein identification. An ion current-based quantification method (IonStar processing pipeline) was described previously (17, 43).

### Bioinformatics Analysis

Ingenuity Pathway Analysis (Qiagen, Redwood City, CA, USA) was used to perform gene ontology enrichment analysis. Hierarchical clustering analysis and heat map visualizations were performed using the *agnes* function in R Package *cluster* and *ggplot2* with the *viridis* color palette, respectively.

### Immunofluorescence

For the Nrf2 nuclear translocation experiment, BMMs were cultured on coverslips and treated with RANKL and M-CSF for 72 h, 5 mM NAC for 16 h, or 50 μM tBHP for 16 h. Coverslips were fixed with 4% paraformaldehyde solution in PBS for 10 minutes at room temperature and permeabilized using 0.1% Triton X-100 for 5 minutes at room temperature. Coverslips were blocked using Image-iT FX signal enhancer (Thermo Fisher Scientific) for 1 hour at room temperature, stained with the primary antibody in 1% BSA/TBST overnight at 4 °C, and stained with the secondary antibody for 1 hour at room temperature. 4,6-diamidino-2-phenylindole (DAPI) (Sigma) was used as a counterstain for nuclei. The coverslips were mounted using ProLong Gold antifade mountant (Thermo) and images were obtained using a fluorescence microscope (Leica, Wetzlar, Germany).

### Western blotting

For experiments studying the Keap1-Nrf2 pathway, cells were cultured in 6-well plates and pre-treated with the indicated concentrations of tBHQ or 25 µM MG −132 for 4 h. For MAPK and NFκB activation experiments, stable-transfected RAW264.7 cells were cultured in 6-well plates and starved in serum-free medium containing 5 mM NAC for 16 h. Cells were subsequently induced with RANKL (200 ng/mL) and M-CSF (100 ng/mL) for the indicated times. Western blotting was performed as described previously (44). The primary antibodies used in this study were as follows: Nrf2 (H-300) and Nrf2 (C-20) (1:100, Santa Cruz Biotechnology, Dallas, TX, USA), Keap1 (E-20) (1:100, SCBT), phospho-p38 (Thr180/Tyr182) (1:1000, Cell Signaling Technology), p38 (1:1000, CST), phospho-ERK1 (Thr202/Tyr204) + ERK2 (Thr186/Tyr187) (1:100, Abcam), ERK1/2 (1:1000, CST), phospho-NFκB p65 (Ser536) (1:1000, CST), NFκB p65 (1:1000, CST), and β-actin (1:4000, SCBT). Densitomety analysis was performed using ImageJ(45) and normalized to the β-actin signal. Relative phosphorylation of was presented as the ratio between the phosphorylated normalized to the nonphosphorylated/total protein. NAC, tBHQ, and tBHP were obtained from Sigma-Aldrich (St. Louis, MO, USA), and MG-132 was obtained from Selleck Chemicals (Houston, TX, USA).

## ACKNOWLEDGMENTS

This work was supported by the National Institute of Arthritis and Musculoskeletal and Skin Diseases (NIAMS, AR061052), the National Institute of Aging (NIA, AG048388) awarded to Dr. Shuying Yang. We would like to thank Dr. Ding Xu and Dr. Sunao Takeshita for their generous gifts of the mRANKL-His (K158-D316) vector and CMG14-12 cell line, respectively.

## Figure 1—source data 1

This Excel sheet contains the micro-CT numerical data and summary statistics represented in Fig. 1E-I.

**Figure 2—figure supplement 1.**
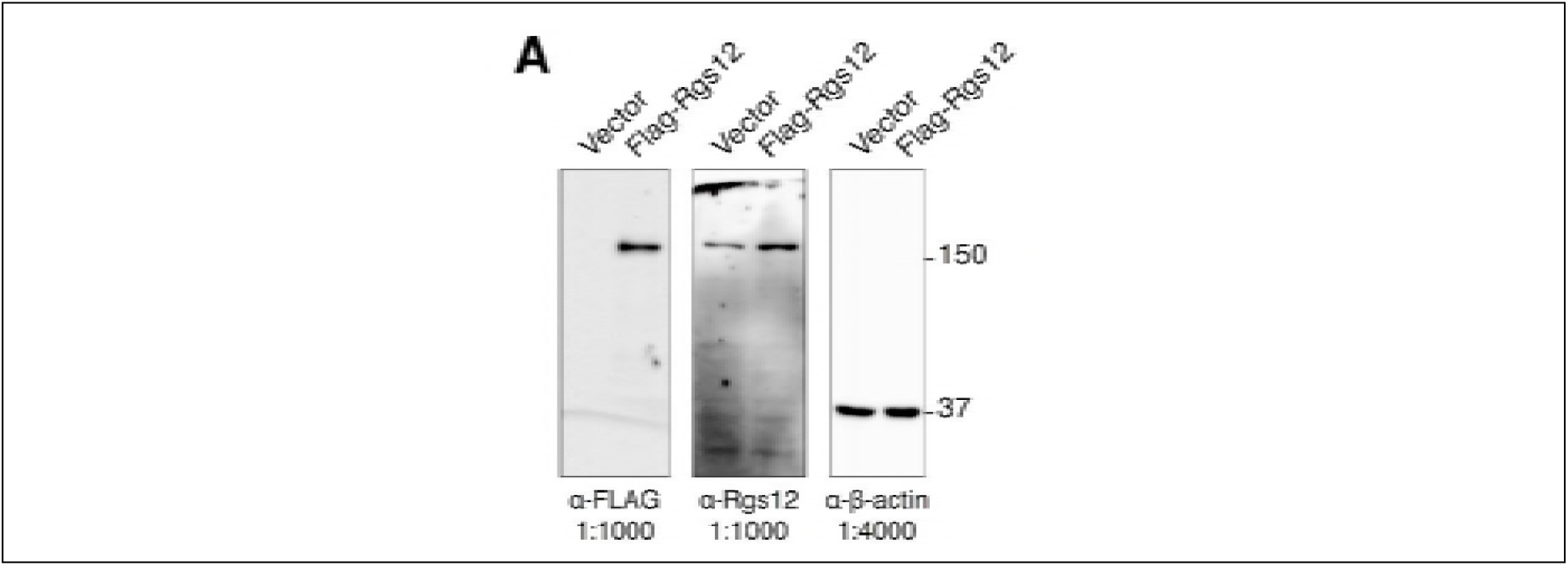
Complete western blots used for Figure 2C.

## Figure 3—source data 1

Table of proteomics data presented in Fig. 3A,B,D,E. Quantitative proteomics analysis of 3,714 proteins in Rgs12 knockout versus wild-type OCs at different timepoints of differentiation. Statistical comparisons between groups were evaluated by Student’s t-test (*P* < 0.05, *n* = 3).

**Figure 5—figure supplement 1.**
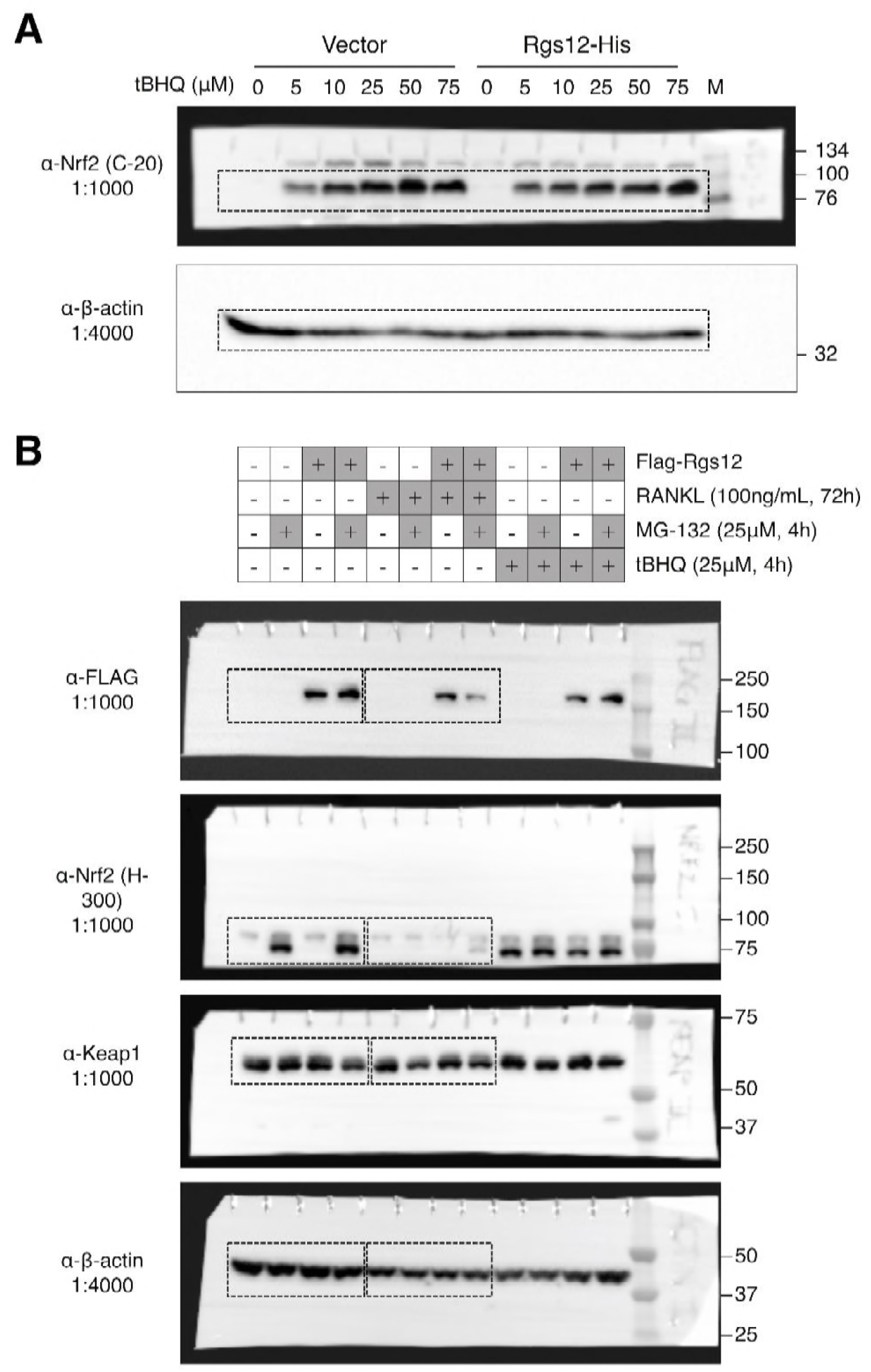
(A) Complete western blots used in Figure 5B. Sections shown in Figure 5B are highlighted with dashed boxes. Transfected RAW264.7 cells were induced with the indicated dosages of tBHQ for 4 h. (B) Complete western blots used in Figure 5D. Sections shown in Figure 5D are highlighted with dashed boxes. RAW264.7 cells stably-transfected with empty vector or Flag-Rgs12 were treated with the following: RANKL (100 ng/ml, 72 h), MG-132 (25 μM, 4 h), and/or tBHQ (25μM,4h). tBHQ, *tert*-butylhydroquinone.

**Figure 6—figure supplement 1.**
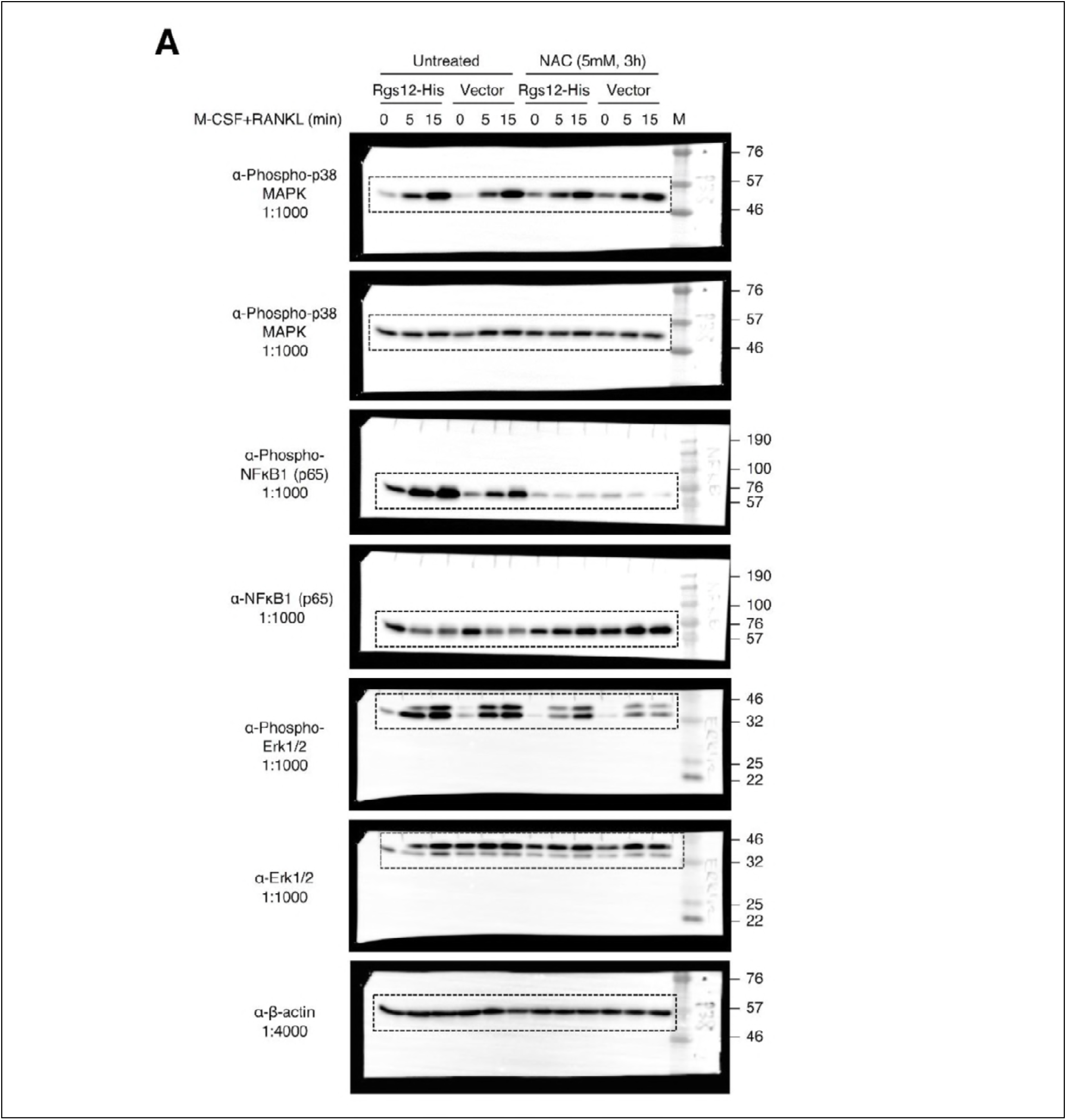
(A) Complete western blots used in Figure 7A. Sections shown in Figure 6C are highlighted with dashed boxes.

